# Nirmatrelvir-resistant SARS-CoV-2 is efficiently transmitted in Syrian hamsters

**DOI:** 10.1101/2022.09.28.509903

**Authors:** Rana Abdelnabi, Dirk Jochmans, Kim Donckers, Bettina Trüeb, Nadine Ebert, Birgit Weynand, Volker Thiel, Johan Neyts

## Abstract

The SARS-CoV-2 main protease (3CLpro) is one of the promising therapeutic target for the treatment of COVID-19. Nirmatrelvir is the only the 3CLpro inhibitor authorized for treatment of COVID-19 patients at high risk of hospitalization; other 3Lpro inhibitors are in development. We recently repored on the *in vitro* selection of a SARS-CoV2 3CLpro (L50F-E166A-L167F; short 3CLpro^res^) virus that is cross-resistant with nirmatrelvir and yet other 3CLpro inhibitors. Here, we demonstrate that the resistant virus replicates efficiently in the lungs of intranassaly infected hamsters and that it causes a lung pathology that is comparable to that caused by the WT virus. Moreover, 3CLpro^res^ infected hamsters transmit the virus efficiently to co-housed non-infected contact hamsters. Fortunately, resistance to Nirmatrelvir does not readily develop (in the clinical setting) since the drug has a relatively high barrier to resistance. Yet, as we demonstrate, in case resistant viruses emerge, they may easily spread and impact therapeutic options for others. Therefore, the use of SARS-CoV-2 3CLpro protease inhibitors in combinations with drugs that have a different mechanism of action, may be considered to avoid the development of drug-resistant viruses in the future.

## Introduction

The Severe acute respiratory syndrome coronavirus 2 (SARS-CoV-2), the causative agent of COVID-19, has had a devastating impact on global public health since its emergence in Wuhan (China) in December 2019. So far, three antiviral drugs have been approved/authorized for clinical use in COVID-19 patients i.e. the nucleoside analoges remdesivir and molnupiravir and the viral protease (3CLpro/Mpro) inhibitor nirmatrelvir *(1)*.

SARS-CoV-2 3CLpro is a cysteine protease which cleaves the two viral polyproteins (pp1a and pp1ab) at eleven different sites, resulting in various non-structural proteins, which are essential for viral replication *(2, 3)*. Interestingly, the cleavage site of the SARS-CoV-2 3CLpro substrate is not recognized by any known human proteases *(4, 5)*. Therefore, 3CLpro is a selective antiviral drug target. Nirmatrelvir (PF-07321332) is a peptidomimetic reversible covalent inhibitor of 3CLpro that is co-formulated with the pharmacokinetic enhancer ritonavir (the resulting combination being marketed as Paxlovid) *(6)*. When treatment is initiated during the first days after symptom onset, it results in roughly 90% protection against severe COVID-19 and hospitalization *(7)*. Besides nirmatrelvir, other 3CLpro inhibitors are currently in clinical development such as the non-peptidic, non-covalent inhibitor, ensitrelvir (S-217622) *(8)*.

The emergence of drug-resistant viruses is a major concern when using antivirals. Development of drug resistance with a subsequent therapeutic failure has been reported during antiviral treatment against different infections including with the human immunodeficiency virus (HIV), hepatitis B virus (HBV), hepatitis C virus (HCV), herpesviruses and influenza viruses *(9, 10)*. Moreover, transmission of drug resistant viruses has also been reported for HIV *(11)* and influenza virus *(12)*. Selection of remdesivir-resistant variants has been reported in cell culture and in the clinical settings *(13–15)*. On the other hand, there are no clear data available yet about emergence of resistant virus variants to molnupiravir or nirmatrelvir in treated patients. Recently, we have reported on *in vitro* selection of a SARS-CoV-2 resistant virus against a first generation 3CLpro inhibitor (ALG-097161) that is cross-resistant to several 3CLpro inhibitors including nirmatrelvir *(16)*. The identified resistant virus carries three amino acid substitutions in the 3CLpro (L50F-E166A-L167F) that result in a more than 20 fold increase in EC_50_ values for different 3CLpro-inhibitors *(16)*. These substitutions are associated with a significant loss of the 3CLpro activity in enzymatic assays, suggesting a subsequent reduction in viral fitness *(16)*.

Here, we aim to (i) explore the infectivity and virulence of the *in vitro* selected 3CLpro (L50F-E166A-L167F) nirmatrelvir resistant (3CLpro^res^) virus in Syrian hamsters and (ii) assess the transmission potential of this *in vitro* selected drug resistant virus from intranasally infected index hamsters to non-infected contact hamsters.

## Results

To assess the infectivity and the transmission potential of the 3CLpro (L50F-E166A-L167F) nirmatrelvir resistant (3CLpro^res^) virus in animals, two groups of index hamsters (each n=12) were intranasally infected with 1×10^4^ TCID_50_ of either the wild-type (WT) SARS-CoV-2 virus (USA-WA1/2020) or the reverse engineered nirmatrelvi resistant virus carrying substitutions L50F-E166A-L167F in the 3CLpro *(16)*.On day 1 post-infection, each of the index hamsters was co-housed in a cage with a contact hamster. The co-housing continued until 3 days after start of contact (Fig. 1A). All hamsters were then euthanized (day 4 pi). Index hamsters infected with the WT virus had median viral RNA and infectious virus loads in the lungs of 1.3×10^7^ genome copies/mg tissue and 1.2×10^5^ TCID_50_/mg tissue, respectively (Fig. 1B/C). The 3CLpro^res^ virus replicated also efficiently in the lungs of infected index hamsters but with lower viral loads than the WT virus [for the 3CLpro^res^ median viral RNA load= 3.1×10^6^ genome copies/mg lung tissue [p=0.0009] and a median infectious virus titers= 1.6×10^3^ TCID_50_/mg lung tissue [p<0.0001]) which is respectively 0.6 and 1.9 log_10_ lower than is the case for WT virus (Fig. 1B/C)].

**Fig. 1.**
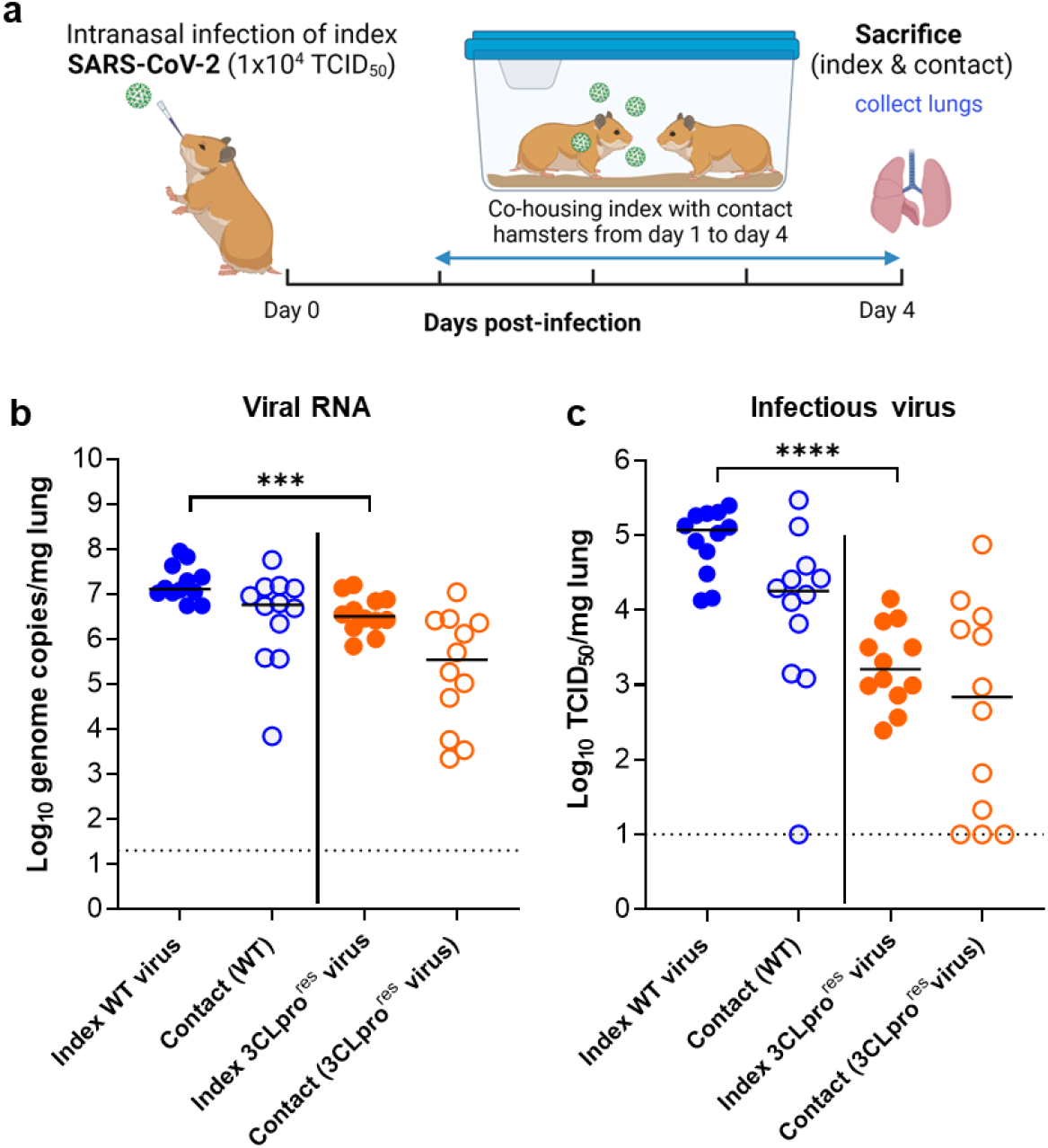
The effect of 3CLpro substitutions, that are associated with nirmatrelvir-resistance, on viral transmission to contact hamsters. (a) Set-up of the study. (b) Viral RNA levels in the lungs of index hamsters infected with 10^4^ TCID_50_ of either the wild-type (WT) SARS-CoV-2 isolate (USA-WA1/2020) or the 3CLpro (L50F-E166A-L167F) nirmatrelvir resistant (3CLpro^res^) virus (closed circles) and contact hamsters (open circles) at day 4 post-infection (pi) are expressed as log_10_ SARS-CoV-2 RNA copies per mg lung tissue. Individual data and median values are presented. (c) Infectious viral loads in the lungs of SARS-CoV-2 WT and (3CLpro^res^) virus -infected index hamsters (closed circles) and contact hamsters (open circles) at day 4 pi are expressed as log_10_ TCID_50_ per mg lung tissue. Individual data and median values are presented. Data were analyzed with the Mann-Whitney U test, ***p<0.001, ****p<0.0001. Data presented are from 2-independent studies with a total n=12 per group. Panel (a) designed by Biorender.

All sentinels that had been co-housed with either WT or the 3CLpro^res^ virus-infected index hamsters had detectable viral RNA in their lungs in the range of (6.9×10^3^-5.7×10^7^) and (2.2×10^3^-1.1×10^7^) genome copies/mg lung tissue, respectively (Fig. 1B). For contacts co-housed with index hamsters that had been infected with WT virus, infectious virus titers were detected in the lungs of 11 out of 12 contact animals with infectious titer loads in the lungs ranging from 1.2×10^3^-2×10^5^ TCID_50_/mg lung tissue (Fig. 1C). On the other hand, 9 out of 12 contacts that had been co-housed with index hamsters infected with the 3CLpro^res^ virus, became infected and infectious virus titers in their lungs ranged between 2×10^1^-7.5×10^4^ TCID_50_/mg lung (Fig. 1C).

Histological study of lungs of the two infected index groups revealed that both the WT and the 3CLpro^res^ virus caused comparable pathological signs including endothelialitis, peri-vascular inflammation, peri-bronchial inflammation and bronchopneumonia (Fig. 2A). Moreover, the median cumulative histopathological lung scores of index hamsters infected with either the WT virus or the 3CLpro^res^ were not different (Fig. 2B).

**Fig. 2.**
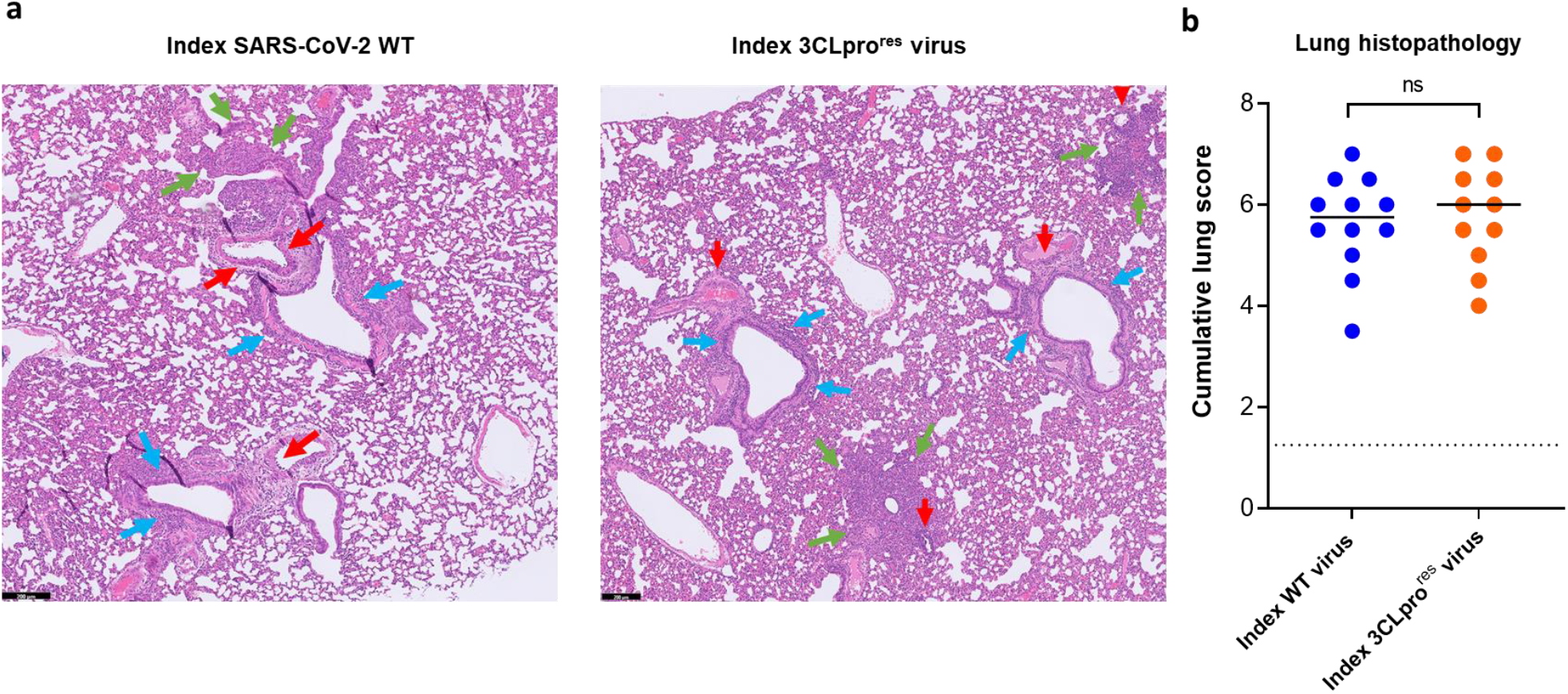
Histopathology of lungs of Syrian hamsters infected with either the wild-type SARS-CoV-2 or the 3CLpro (L50F-E166A-L167F) nirmatrelvir resistant virus. (a) Representative H&E images of lungs of hamsters infected with 10^4^ TCID_50_ of either the wild-type (WT) SARS-CoV-2 virus (USA-WA1/2020) or the 3CLpro (L50F-E166A-L167F) nirmatrelvir resistant (3CLpro^res^) virus at day 4 post-infection (pi) showing peribronchial inflammation (blue arrows), peri-vascular inflammation with vasculitis (red arrows), and foci of bronchopneumonia (green arrows). Scale bars, 200 μm. (b) Cumulative severity score from H&E stained slides of lungs from hamsters infected with the WT virus or the (3CLpro^res^) virus at day 4 pi. Individual data and median values are presented and the dotted line represents the median score of untreated non-infected hamsters. Data were analyzed with the Mann-Whitney U test. Data presented are from 2-independent studies with a total n=12 per group.

Deep sequencing analysis of viral RNA isolated from the lungs of index and contact hamsters infected with the 3CLpro^res^ virus revealed that the L50F-E166A-L167F subsstitutions were maintained in the 3CLpro of this resistant virus after replication in hamsters, indicating the genomic stability of these mutations.

## Discussion

Emergence of drug-resistant variants is a major problem during antiviral treatment for several acute and chronic viral infection such as HIV, HCV and influenza virus infections. So far SARs-CoV-2 drug resistant variants have been reported only for the treatment with the polymerase inhibitor remdesivir *(14)*. Recently, we have reported on the *in vitro* selection of a resistant virus against 3CLpro inhibitors including nirmatrelvir and ensetrilvir, that carries the L50F-E166A-L167F substitutions in the 3CLpro protein *(16)*. In a FRET-based assay, the enzymatic activities of recombinant 3CLpro proteins carrying the L50F, E166A and L167F subsstitutions alone or combined proved to be significantly lower, compared to that of the WT protein *(16)*. In another similar study, *in vitro* selection of resistant variants against nirmatrelvir resulted in identification of several resistance-associated subsstitutions with the E166V subsstitution resulting in the strongest resistance phenotype *(17)*. However, this subsstitution at position 166 resulted in loss of the virus replication fitness *in vitro* that was restored by compensatory subsstitutions such as L50F and T21I *(17)*.

Here, we assed the *in vivo* fitness of of this 3CLpro (L50F-E166A-L167F) nirmatrelvir (and other protease inhibitors) resistant virus in terms of infectivity and transmission potential in a hamster infection model. Intranasal infection of Syrian hamsters with either the WT or the 3CLpro^res^ variant resulted in efficient replication of the virus in the lungs. However, the viral RNA loads and infectious titers in the lungs of hamsters infected with the 3CLpro^res^ virus were 0.6 and 1.9 log_10_ lower compared to the WT virus-infected index hamsters. On the other hand, both the WT and the 3CLpro^res^ virus caused a comparable lung pathology in the infected index hamsters.

Co-housing of each index animal, infected with either the WT or the 3CLpro^res^ virus, in a cage with a contact (non-infected hamster) for 3 days revealed efficient transmission of either virus between infected and non-infected hamsters. All contact hamsters from both groups had detectable viral RNA loads in their lungs. Infectious virus titers were detected in the lungs of respectively 92% and 75% of the sentinels co-housed with animals that had been infected with either the WT virus or 3CLpro^res^ virus. The discrepancy between the number of contact hamsters with viral RNA in the lungs and detectable infectious virus in their lungs might be explained by the fact that all the contact animals were sacrificed at the same day (day 3 post first contact) whereas transmission most likely does not occur in a synchronized way; some may possibly soon, others later after contact be infected. Accordingly, for animals that acquired the virus rather late after the first contact, viral RNA might already be detectable in the lungs, but one or two more days may have been required to allow detection of infectious titers.

Taken together, these results show that the 3CLpro (L50F-E166A-L167F) nirmatrelvir resistant virus is able to efficiently replicate in the lungs of Syrian hamsters and that this variant can be transmitted via direct contact to co-housed naive hamsters. Two of the selected substitutions (i.e. L50F and L167F) have been reported to be detected (at low frequencies) in SARS-CoV-2 viruses naturally circulating in the population (17). Moreover, the Paxlovid label indicates that the E166V subsititution is among the treatment-emergent substitution, which is more common in nirmatrelvir/ritonavir treated subjects relative to placebo-treated subjects. These observations suggest that the *in vitro* selected 3CLpro (L50F-E166A-L167F) resistant virus may also emerge in treated patients with increased use of Paxlovid or other 3CLpro targeting drugs in the future. The emergence of drug-resistant SARS-CoV-2 variants that can be efficiently transmitted in the population is of serious concern. It may result in a loss of important therapeutic option for patients that acquire the drug-resistant virus. Efficient transmission of the anti-influenza drugs amantadine and rimantadine has been well documented *(12)* although it should be mentioned that the barrier to resistance of these amantadanes is much lower than that of the SARS-CoV2 3CLpro inhibitors currently being developed. For the treatment of infections with HIV and HCV fixed-dose combinations prevent the emergence of resistant variants *(18)*. It will be prudent and important to explore the efficacy and impact on potential resistance development of combinations of SARS-CoV-2 3CLpro inhibitors with drugs with a non-overlapping resistance profile.

## Methods

### SARS-CoV-2

The WT SARS-CoV-2 strain used in this study, SARS-CoV-2 USA-WA1/2020 (EPI_ISL_404895), was obtained through BEI Resources (ATCC) cat. No. NR-52281, batch 70036318. This isolate has a close relation with the prototypic Wuhan-Hu-1 2019-nCoV (GenBank accession 112 number MN908947.3) strain as confirmed by phylogenetic analysis. The reverse engineered 3CLpro (L50F-E166A-L167F) nirmatrelvir resistant (3CLpro^res^) virus used here has been described before (16). Live virus-related work was conducted in the high-containment A3 and BSL3+ facilities of the KU Leuven Rega Institute (3CAPS) under licenses AMV 30112018 SBB 219 2018 0892 and AMV 23102017 SBB 219 20170589 according to institutional guidelines.

### Cells

Vero E6 cells (African green monkey kidney, ATCC CRL-1586) were cultured in minimal essential medium (MEM, Gibco) supplemented with 10% fetal bovine serum (Integro), 1% non-essential amino acids (NEAA, Gibco), 1% L-glutamine (Gibco) and 1% bicarbonate (Gibco). End-point titrations on Vero E6 cells were performed with medium containing 2% fetal bovine serum instead of 10%. Cells were kept in a humidified 5% CO_2_ incubator at 37°C.

### Infection and transmission study

The hamster infection model of SARS-CoV-2 has been described before *(19, 20)*. Female Syrian hamsters (Mesocricetus auratus) were purchased from Janvier Laboratories and kept per two in individually ventilated isolator cages (IsoCage N Bio-containment System, Tecniplast) at 21°C, 55% humidity and 12:12 day/night cycles. Housing conditions and experimental procedures were approved by the ethics committee of animal experimentation of KU Leuven (license P065-2020). For infection, two groups of index female hamsters of 6-8 weeks old were anesthetized with ketamine/xylazine/atropine and inoculated intranasally with 50 μL containing 1×10^4^ TCID_50_ of either SARS-CoV-2 WT virus or the 3CLpro^res^ virus(day 0). In the morning of day1 post-infection (pi), each index hamster was co-housed with a contact (sentinel) hamster (non-infected hamsters) in one cage and the co-housing continued until the sacrifice day i.e. 3 days after start of exposure. On day 4 pi, animals were euthanized by intraperitoneal injection of 500 μL Dolethal (200 mg/mL sodium pentobarbital) and lungs were collected for further analysis. Two independent studies were performed each with n=6 per group.

### SARS-CoV-2 RT-qPCR

Hamster lung tissues were collected after sacrifice and were homogenized using bead disruption (Precellys) in 350 μL TRK lysis buffer (E.Z.N.A.^®^ Total RNA Kit, Omega Bio-tek) and centrifuged (10.000 rpm, 5 min) to pellet the cell debris. RNA was extracted according to the manufacturer’s instructions. RT-qPCR was performed on a LightCycler96 platform (Roche) using the iTaq Universal Probes One-Step RT-qPCR kit (BioRad) with N2 primers and probes targeting the nucleocapsid *(20)*. Standards of SARS-CoV-2 cDNA (IDT) were used to express viral genome copies per mg tissue *(19)*.

### End-point virus titrations

Lung tissues were homogenized using bead disruption (Precellys) in 350 μL minimal essential medium and centrifuged (10,000 rpm, 5min, 4°C) to pellet the cell debris. To quantify infectious SARS-CoV-2 particles, endpoint titrations were performed on confluent Vero E6 cells in 96-well plates. Viral titers were calculated by the Reed and Muench method *(21)* using the Lindenbach calculator and were expressed as 50% tissue culture infectious dose (TCID_50_) per mg tissue.

### Histology

For histological examination, the lungs were fixed overnight in 4% formaldehyde and embedded in paraffin. Tissue sections (5 μm) were analyzed after staining with hematoxylin and eosin and scored blindly for lung damage by an expert pathologist. The scored parameters, to which a cumulative score of 1 to 3 was attributed, were the following: congestion, intra-alveolar hemorrhagic, apoptotic bodies in bronchus wall, necrotizing bronchiolitis, perivascular edema, bronchopneumonia, perivascular inflammation, peribronchial inflammation and vasculitis.

### Deep sequencing and analysis of whole genome sequences

The viral RNAs isolated from the lungs of infected animals were subjected to deep sequencing analysis of the full genome sequence through ARTIC SARS-CoV-2 RNA-Seq service provided by Eurofins Genomics. Alignment of the obtained sequences was performed using the Geneious Prime software.

### Statistics

The detailed statistical comparisons, the number of animals and independent experiments that were performed is indicated in the legends to figures. “Independednt experiments” means that experiments were repeated separately on different days. The analysis of histopathology was done blindly. All statistical analyses were performed using GraphPad Prism 9 software (GraphPad, San Diego, CA, USA). Statistical significance was determined using the non-parametric Mann Whitney U-test. P-values of <0.05 were considered significant.

### Ethics

Housing conditions and experimental procedures were done with the approval and under the guidelines of the ethics committee of animal experimentation of KU Leuven (license P065-2020).

## Acknowledgments

We thank Carolien De Keyzer, Lindsey Bervoets, Thibault Francken, Stijn Hendrickx and Niels Cremers for excellent technical assistance. We thank Prof. Jef Arnout and Dr. Annelies Sterckx (KU Leuven Faculty of Medicine, Biomedical Sciences Group Management) and Animalia and Biosafety Departments of KU Leuven for facilitating the animal studies.

## Author Contributions

R.A. and J.N. designed the studies; R.A and B.W. performed the studies and analyzed data; R.A. made the graphs; B.W., D.J. and J.N. provided advice on the interpretation of data; R.A. and J.N. wrote the paper; B.T., N.E.J and V.T. provided the revesrse engineered virus variant; R.A., V.T. and J.N. supervised the study; D.J. and J.N. acquired funding.

## References

1. A. C. Puhl, T. R. Lane, F. Urbina, S. Ekins, The Need for Speed and Efficiency: A Brief Review of Small Molecule Antivirals for COVID-19. Front. Drug Discov. 0, 3 (2022).

2. Z. Jin, X. Du, Y. Xu, Y. Deng, M. Liu, Y. Zhao, B. Zhang, X. Li, L. Zhang, C. Peng, Y. Duan, J. Yu, L. Wang, K. Yang, F. Liu, R. Jiang, X. Yang, T. You, X. Liu, X. Yang, F. Bai, H. Liu, X. Liu, L. W. Guddat, W. Xu, G. Xiao, C. Qin, Z. Shi, H. Jiang, Z. Rao, H. Yang, Structure of Mpro from SARS-CoV-2 and discovery of its inhibitors. Nature 582, 289–293 (2020).

3. T. Pillaiyar, M. Manickam, V. Namasivayam, Y. Hayashi, S. H. Jung, An overview of severe acute respiratory syndrome-coronavirus (SARS-CoV) 3CL protease inhibitors: Peptidomimetics and small molecule chemotherapy J Med. Chem. 59, 6595–6628 (2016).

4. K. Fan, L. Ma, X. Han, H. Liang, P. Wei, Y. Liu, L. Lai, The substrate specificity of SARS coronavirus 3C-like proteinase. Biochem.Biophys. Res. Commun. 329, 934–940 (2005).

5. S. Ullrich, C. Nitsche, The SARS-CoV-2 main protease as drug *targetBioorganic Med*. Chem. Lett. 30 (2020), doi:10.1016/j.bmcl.2020.127377.

6. D. R. Owen, C. M. N. Allerton, A. S. Anderson, L. Aschenbrenner, M. Avery, S. Berritt, B. Boras, R. D. Cardin, A. Carlo, K. J. Coffman, A. Dantonio, L. Di, H. Eng, R. Ferre, K. S. Gajiwala, S. A. Gibson, S. E. Greasley, B. L. Hurst, E. P. Kadar, A. S. Kalgutkar, J. C. Lee, J. Lee, W. Liu, S. W. Mason, S. Noell, J. J. Novak, R. S. Obach, K. Ogilvie, N. C. Patel, M. Pettersson, D. K. Rai, M. R. Reese, M. F. Sammons, J. G. Sathish, R. S. P. Singh, C. M. Steppan, A. E. Stewart, J. B. Tuttle, L. Updyke, P. R. Verhoest, L. Wei, Q. Yang, Y. Zhu, An Oral SARS-CoV-2 Mpro Inhibitor Clinical Candidate for the Treatment of COVID-19. Science (80-.)., eabl4784 (2021).

7. E. Mahase, Covid-19: Pfizer’s paxlovid is 89% effective in patients at risk of serious illness, company reports. BMJ 375, n2713 (2021).

8. J. D. A. Tyndall, S-217622, a 3CL Protease Inhibitor and Clinical Candidate for SARS-CoV-2. J. Med. Chem. 65, 6496–6498 (2022).

9. M. G. Ison, F. G. Hayden, A. J. Hay, L. V Gubareva, E. A. Govorkova, E. Takashita, J. L. McKimm-Breschkin, Influenza polymerase inhibitor resistance: Assessment of the current state of the art - A report of the isirv Antiviral group. Antiviral Res. 194, 105158 (2021).

10. L. Menéndez-Arias, R. Delgado, Update and latest advances in antiretroviral therapy. Trends Pharmacol. Sci. 43, 16–29 (2022).

11. C. Guo, Y. Wu, Y. Zhang, X. Liu, A. Li, M. Gao, T. Zhang, H. Wu, G. Chen, X. Huang, Transmitted Drug Resistance in Antiretroviral Therapy-Naive Persons With Acute/Early/Primary HIV Infection: A Systematic Review and Meta-Analysis. Front. Pharmacol. 12 (2021), doi: 10.3389/fphar.2021.718763.

12. W. He, W. Zhang, H. Yan, H. Xu, Y. Xie, Q. Wu, C. Wang, G. Dong, Distribution and evolution of H1N1 influenza A viruses with adamantanes-resistant mutations worldwide from 1918 to 2019. J. Med. Virol. 93, 3473–3483 (2021).

13. D. Focosi, F. Maggi, S. McConnell, A. Casadevall, Very low levels of remdesivir resistance in SARS-COV-2 genomes after 18 months of massive usage during the COVID19 pandemic: A GISAID exploratory analysis. Antiviral Res. 198, 105247 (2022).

14. S. Gandhi, J. Klein, A. J. Robertson, M. A. Peña-Hernández, M. J. Lin, P. Roychoudhury, P. Lu, J. Fournier, D. Ferguson, S. A. K. Mohamed Bakhash, M. Catherine Muenker, A. Srivathsan, E. A. Wunder, N. Kerantzas, W. Wang, B. Lindenbach, A. Pyle, C. B. Wilen, O. Ogbuagu, A. L. Greninger, A. Iwasaki, W. L. Schulz, A. I. Ko, De novo emergence of a remdesivir resistance mutation during treatment of persistent SARS-CoV-2 infection in an immunocompromised patient: a case report. Nat. Commun. 13, 1547 (2022).

15. L. J. Stevens, A. J. Pruijssers, H. W. Lee, C. J. Gordon, E. P. Tchesnokov, J. Gribble, A. S. George, T. M. Hughes, X. Lu, J. Li, J. K. Perry, D. P. Porter, T. Cihlar, T. P. Sheahan, R. S. Baric, M. Götte, M. R. Denison, Mutations in the SARS-CoV-2 RNA-dependent RNA polymerase confer resistance to remdesivir by distinct mechanisms. Sci. Transl. Med. 14, eabo0718 (2022).

16. D. Jochmans, C. Liu, K. Donckers, A. Stoycheva, S. Boland, S. K. Stevens, C. De Vita, B. Vanmechelen, P. Maes, B. Trüeb, N. Ebert, V. Thiel, S. De Jonghe, L. Vangeel, D. Bardiot, A. Jekle, L. M. Blatt, L. Beigelman, J. A. Symons, P. Raboisson, P. Chaltin, A. Marchand, J. Neyts, J. Deval, K. Vandyck, The substitutions L50F, E166A and L167F in SARS-CoV-2 3CLpro are selected by a protease inhibitor in vitro and confer resistance to nirmatrelvir. bioRxiv, 2022.06.07.495116 (2022).

17. S. Iketani, H. Mohri, B. Culbertson, S. J. Hong, Y. Duan, M. I. Luck, M. K. Annavajhala, Y. Guo, Z. Sheng, A.-C. Uhlemann, S. P. Goff, Y. Sabo, H. Yang, A. Chavez, D. D. Ho, Multiple pathways for SARS-CoV-2 resistance to nirmatrelvir. bioRxiv, 2022.08.07.499047 (2022).

18. Z. A. Shyr, Y. S. Cheng, D. C. Lo, W. Zheng, Drug combination therapy for emerging viral diseases. Drug Discov. Today 26, 2367–2376 (2021).

19. S. J. F. Kaptein, S. Jacobs, L. Langendries, L. Seldeslachts, S. ter Horst, L. Liesenborghs, B. Hens, V. Vergote, E. Heylen, K. Barthelemy, E. Maas, C. de Keyzer, L. Bervoets, J. Rymenants, T. van Buyten, X. Zhang, R. Abdelnabi, J. Pang, R. Williams, H. J. Thibaut, K. Dallmeier, R. Boudewijns, J. Wouters, P. Augustijns, N. Verougstraete, C. Cawthorne, J. Breuer, C. Solas, B. Weynand, P. Annaert, I. Spriet, G. Vande Velde, J. Neyts, J. Rocha-Pereira, L. Delang, Favipiravir at high doses has potent antiviral activity in SARS-CoV-2-infected hamsters, whereas hydroxychloroquine lacks activity. Proc. Natl. Acad. Sci. U. S. A. 117, 26955–26965 (2020).

20. R. Boudewijns, H. J. Thibaut, S. J. F. Kaptein, R. Li, V. Vergote, L. Seldeslachts, J. Van Weyenbergh, C. De Keyzer, L. Bervoets, S. Sharma, L. Liesenborghs, J. Ma, S. Jansen, D. Van Looveren, T. Vercruysse, X. Wang, D. Jochmans, E. Martens, K. Roose, D. De Vlieger, B. Schepens, T. Van Buyten, S. Jacobs, Y. Liu, J. Martí-Carreras, B. Vanmechelen, T. Wawina-Bokalanga, L. Delang, J. Rocha-Pereira, L. Coelmont, W. Chiu, P. Leyssen, E. Heylen, D. Schols, L. Wang, L. Close, J. Matthijnssens, M. Van Ranst, V. Compernolle, G. Schramm, K. Van Laere, X. Saelens, N. Callewaert, G. Opdenakker, P. Maes, B. Weynand, C. Cawthorne, G. Vande Velde, Z. Wang, J. Neyts, K. Dallmeier, STAT2 signaling restricts viral dissemination but drives severe pneumonia in SARS-CoV-2 infected hamsters. Nat. Commun. 11, 5838 (2020).

21. L. J. Reed, H. Muench, A simple method of estimating fifty percent endpoints. Am. J. Hyg. 27, 493–497 (1938).

